# AI-Based Comparison of UniProt annotations and Literature Across Human Kinases

**DOI:** 10.64898/2026.09.23.753935

**Authors:** Jimin Pei, Qian Cong, Jing Zhang, Nick V. Grishin

## Abstract

Curated protein records and their supporting publications organize biological evidence at different levels of detail. Here, we used AI to independently answer a set of biological questions using either UniProt records or the full texts of the literature cited by those records, and then used AI-based evaluation to compare the resulting answers in terms of information coverage and semantic agreement. Fifty human proteins were selected from ten groups in a kinase sequence collection. Target-identity filtering retained 1,749 protein–publication records representing 1,655 distinct publications. For each protein, an AI language model generated one answer set from UniProt and another from the associated literature. A separate AI-based semantic evaluation compared all 2,500 pairwise combinations of the 50 UniProt-derived and 50 literature-derived answer sets, producing 87,500 question-level comparisons. Literature provided answers to 1,211 of 1,750 protein–question combinations, compared with 1,151 for UniProt. After correction for multiple testing across the 35 questions, literature coverage was significantly higher for autoinhibition and inhibitor type. Mean AI-assigned semantic agreement scores were higher for matched protein pairs (0.692) than for unmatched proteins within the same kinase group (0.384) and for proteins from different kinase groups (0.279). The matching literature-derived answer set ranked first for 49 of the 50 UniProt-derived queries, even after questions about gene symbol and kinase family were excluded. The sole exception was GUCY2C, whose UniProt-derived answer set showed slightly greater agreement with the NPR2 literature-derived answer set than with the GUCY2C literature-derived set. TYR proteins had the greatest overall answer coverage, whereas CK1 and RGC proteins had comparatively limited coverage in UniProt and the literature, respectively. Overall, AI analysis showed that UniProt records and their cited publications contain complementary yet strongly protein-specific biological information. Reporting gaps, question scope, and score aggregation influenced apparent agreement, underscoring that the framework evaluates consistency between AI-derived answers, not their biological accuracy.

## Introduction

Knowledge of protein function comes from both curated database annotations and the experimental literature on which they draw. Studying a protein kinase, for example, often requires information about its catalytic activity, regulation, binding partners, cellular localization, and involvement in disease [1]. The level of detail needed depends on the question being asked. Database entries bring together findings from multiple studies, whereas individual publications provide experimental details that can be essential for interpreting a result, such as the protein construct, mutation, tissue, or treatment condition examined [2–4].

The UniProt database provides protein sequences together with functional annotations assembled through expert curation and computational approaches [2]. Its reviewed records integrate evidence into a common framework that supports biological interpretation and computational analysis. The database also identifies supporting publications, providing a practical connection between protein annotations and their underlying literature. However, the existence of a citation does not establish that every observation in the cited article is represented in the protein record. Conversely, an experimental observation extracted from one publication does not necessarily have the breadth or evidential status of a curated annotation.

Protein kinases are well suited to examining these relationships. Members of the human kinome share related kinase domains but differ widely in their regulatory mechanisms and biological functions. Their classification provides a framework for comparative studies of cell signaling [1]. A comparative question panel can therefore include both properties shared by many kinases and features that distinguish individual proteins. A useful comparison must retain these biological distinctions while recognizing equivalent descriptions expressed in different terminology.

AI language models can organize information from scientific text into structured evidence and readable answers. Previous studies have shown that these models can extract entities and their relationships from scientific publications, capturing information beyond individual keywords [5, 6]. These capabilities allow information from different sources to be compared using a common set of biological questions. However, the answers reflect the model’s interpretation of the supplied evidence. During extraction and synthesis, models may omit relevant findings, misinterpret ambiguous terms, or lose qualifications tied to specific experimental conditions. A clearly written answer is therefore not necessarily an accurate one [7, 8].

Model evaluation introduces an additional layer of interpretation. Semantic comparison can recognize equivalent claims despite differences in wording, but it must distinguish contradictory answers from differences in coverage. When neither source reports a property, assigning a full agreement score would reward shared missingness. When one source provides additional information, a lower agreement score may reflect complementary coverage rather than an erroneous biological claim. Studies of language models used as evaluators have also identified sensitivity to answer presentation and other biases, motivating explicit evaluation rules and inspection of difficult cases [9–11].

Several benchmarks have been developed to evaluate the accuracy of computational methods in extracting information from biomedical literature. PubMedQA draws its questions from article titles and uses abstract conclusions as reference answers, with human annotators assigning yes, no, or maybe labels to a set of questions. [12]. SciFact uses scientific claims derived from citation statements, with human annotators determining whether relevant abstracts support or refute each claim and identifying the sentences that provide the evidence. [13]. In BioASQ, biomedical experts develop questions, identify relevant articles and passages, and write reference answers [14]. Although these benchmarks differ in scope, each relies on human annotation or expert curation to provide a basis for evaluating computational results.

Other benchmarks examine a wider range of evidence sources and research tasks. QASPER evaluates question answering over full-text natural-language-processing papers [15]. QASPER questions were written by readers who saw only the title and abstract, while separate annotators prepared reference answers and identified supporting passages, figures, or tables from the complete paper [15]. LAB-Bench evaluates practical biological research skills through multiple-choice questions covering literature retrieval, figure and table interpretation, database access, protocol troubleshooting, sequence analysis, and molecular cloning [16].

LAB-Bench questions and reference answers were developed specifically for the benchmark through expert writing and review or generated programmatically from biological databases and sequence calculations [16]. SciAssess evaluates scientific knowledge, document comprehension, and reasoning across biology, chemistry, materials science, and medicine, using tasks involving text, figures, tables, and molecular structures [17]. ScholarQABench combines existing datasets with expert-written questions that require synthesizing findings from multiple papers [18]. The accompanying OpenScholar system retrieves and ranks relevant passages and can gather additional evidence while revising its answers [18].

The sources of reference information also vary across benchmarks. In the examples above, questions, labels, scoring rubrics, and reference answers were generally created or adapted specifically for evaluation, with contributors ranging from supervised trainees to experienced domain specialists and with different procedures for checking their work. Some earlier benchmarks also drew on established database annotations. For example, BioCreative Task 2 [19] linked full-text articles to existing, manually curated protein–Gene Ontology annotations from GOA for its training data. GOA curators provided the test annotations, and three curators assessed the proteins, Gene Ontology terms, and supporting passages identified by participating systems. BioCreative thus combined existing database annotations with expert curation and assessment carried out specifically for the benchmark.

These benchmarks have focused on evaluating how accurately AI systems interpret scientific literature, retrieve supporting evidence, and produce answers or structured information consistent with reference annotations or expert judgments. Less attention has been given to whether AI can systematically capture the range of relevant biological information spread across a defined collection of publications. The present study addresses this complementary aspect by using pre-existing UniProt annotations as a standardized evidence source and comparing AI-generated answers from UniProt with independently generated answers from the full texts of their cited publications. We evaluate information coverage and semantic agreement using a common set of questions about biological properties, and test whether the answer profiles preserve features specific to individual proteins and kinase groups. Questions about substrates, regulation, localization, and other properties may have several valid answers that depend on the biological context and experimental conditions. The framework therefore allows each answer to include multiple relevant findings rather than a single predetermined result.

Here we studied 50 human protein kinases from 10 kinase groups, choosing the five proteins in each group with the most publications cited in UniProt. For each protein, we used AI to independently answer the same biological questions from two sources: its UniProt record and information extracted from the full texts of publications cited in that record. We then used AI to compare the answers for semantic agreement. The analysis addressed three related questions. We first compared answer coverage between the two sources across biological questions and selected proteins. We then tested whether AI-assigned semantic agreement scores could distinguish the matching protein from other proteins. Finally, we assessed whether the AI-derived answer profiles reflected the kinase group assignments. Because the publications were drawn from UniProt reference lists, the two sources represent overlapping bodies of evidence. This design allows us to examine which information is shared, added, or missing when the same protein is described using curated annotations or AI-generated summaries of the cited literature.

## Results

### Study collection and evidence retention

The analysis included 50 human proteins, with five representatives from each of ten kinase groups labeled AGC, CAMK, CK1, CMGC, NEK, OTHER, RGC, STE, TKL, and TYR [20]. The panel included conventional protein kinases and the five RGC proteins (GUCY2C, GUCY2D, GUCY2F, NPR1, and NPR2) with inactive kinase domains. The starting literature collection comprised 2,199 protein–publication records representing 1,790 distinct PubMed identifiers. All recorded PubMed identifiers were found among the citations in their corresponding local UniProt records. Target identity filtering retained 1,587 records in which the target had a primary or co-primary role and 162 records in which the target was identified in a contextual role. The resulting evidence collection contained 1,749 records, or 79.5% of the starting collection, representing 1,655 distinct publications.

The final dataset contained 50 UniProt answer sets and 50 literature answer sets, each with entries for all 35 biological questions. Their exhaustive comparison produced 2,500 protein pairs and 87,500 question comparisons. Of these comparisons, 69,997 received a numeric score and 17,503 were classified as not comparable. Numeric judgments comprised 10,649 semantic matches, 15,244 partial matches, 23,129 mismatches, and 20,975 coverage mismatches (one answer is reported and the other answer is not reported).

### Literature and UniProt provided overlapping but complementary coverage

Across the 1,750 possible protein–question combinations in each source, UniProt provided 1,151 reported answers (65.8%), and literature provided 1,211 (69.2%). Both sources reported an answer for 1,069 combinations, whereas neither reported an answer for 457. Literature alone reported answers for 142 combinations, and UniProt alone did so for 82. Taking the union of their reporting states therefore covered 1,293 combinations (73.9%). This union describes potential coverage; conflicting claims were not reconciled into a new combined answer set.

The net literature advantage was 3.43 percentage points. A bootstrap that resampled proteins within their assigned groups, keeping all questions and both sources together, gave a 95% confidence interval of 1.71–5.09 percentage points. An exploratory permutation test that swapped source labels jointly within each group gave a two-sided P value of 0.0137. The gain was therefore modest in absolute terms but consistent with a systematic difference in coverage within this selected panel.

Coverage varied substantially across questions (Figure 1; Supplementary Table S1). Gene symbols, substrates, and subcellular localizations were reported for all 50 proteins by both sources. Cellular consequences, modification types, and upstream regulators were each reported in 99 of the 100 source specific answer sets. Downstream pathways and activation mechanisms were also broadly covered, with 98 and 96 reported answers, respectively. Broad coverage did not necessarily imply identical content: substrate lists and localization descriptions frequently received partial rather than complete semantic matches.

**Figure 1.**
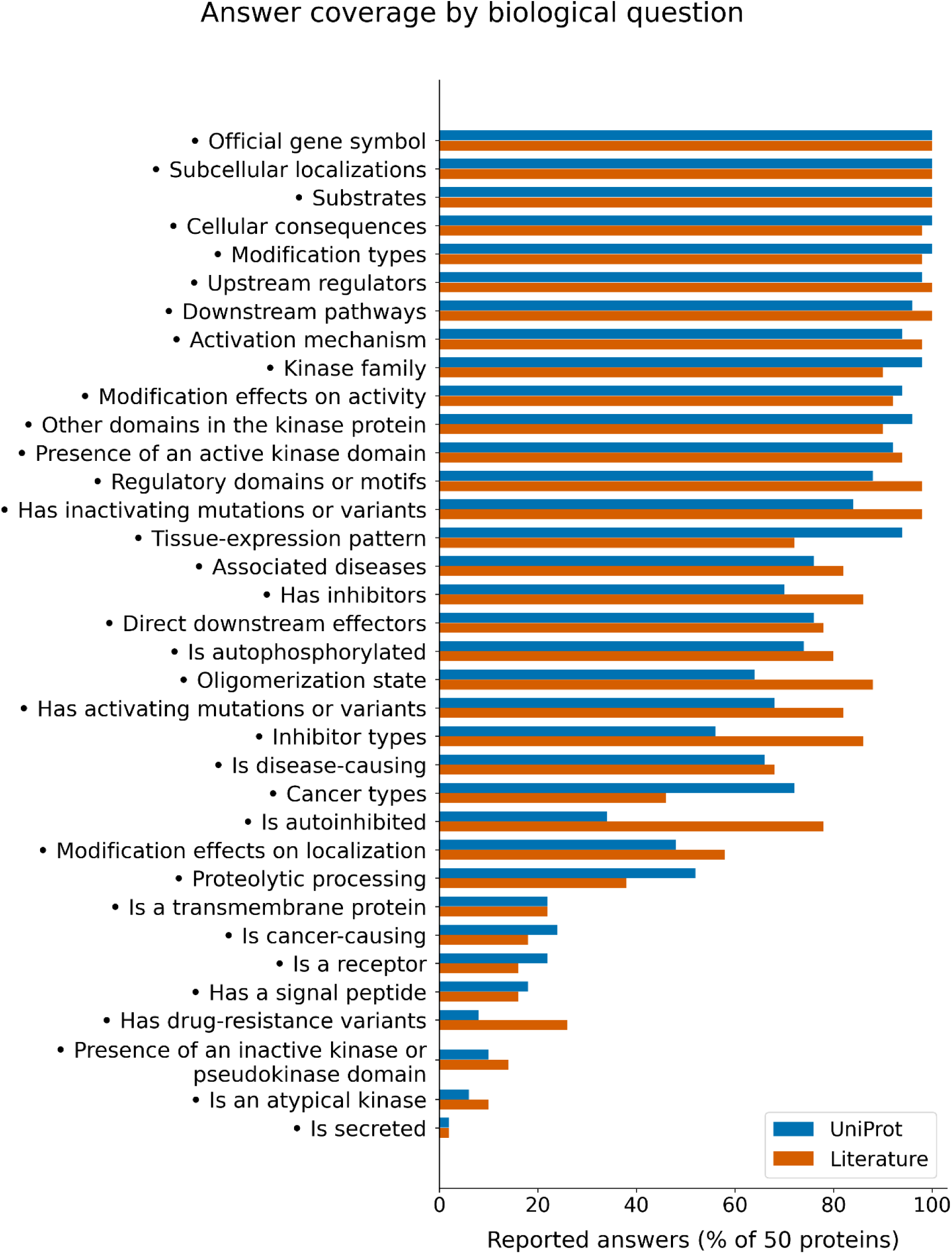
Coverage of the 35 biological questions. Bars show the proportion of the 50 proteins with a reported answer in each source. Questions are ordered by combined coverage. Reporting states are those saved during answer generation, before semantic adjudication. Low coverage indicates limited explicit reporting under the extraction procedure and does not establish that the corresponding property is absent.

At the other end of the distribution, secretion was reported in only two source specific answer sets, both concerning EGFR. Atypical kinase status and the presence of inactive kinase or pseudokinase domains were reported in 8 and 12 answer sets, respectively. Drug resistance variants and signal peptides were each reported in 17, receptor status in 19, and transmembrane status in 22. These counts quantify explicit reporting under the extraction procedure. They do not estimate the prevalence of the corresponding biological properties, because an unreported property was not treated as a negative finding.

The largest literature advantages occurred for autoinhibition, reported for 39 proteins in literature and 17 in UniProt, and inhibitor types, reported for 43 and 28 proteins, respectively. Exact paired tests followed by Holm correction across all 35 questions retained these two differences at the 0.05 level, with adjusted P values of 0.000104 and 0.00208. Literature also had greater observed coverage for oligomerization, drug resistance variants, inhibitors, and activating or inactivating variants, but these differences did not pass the same correction. UniProt had greater observed coverage for cancer types (36 versus 23 proteins), tissue expression (47 versus 36), and proteolytic processing (26 versus 19). Their adjusted P values were 0.0775, 0.222, and 1.000, respectively. These directional differences are informative descriptively, but the present analysis does not establish statistical significance for each of them.

### Coverage and consistency differed across individual proteins

EGFR had the greatest coverage in both sources, with 33 reported answers out of 35. UniProt coverage was also high for INSR and RET, each with 30 reported answers, and TGFBR1, with 29. Literature coverage reached 32 for RET, 31 for TGFBR1, and 30 for INSR. The least covered UniProt protein was VRK2, with 15 reported answers, followed by NEK1 and CSNK1A1, each with 16. The least covered literature protein was GUCY2F, with 12, followed by CSNK1A1 with 16 and CDK1 with 17. GUCY2F had only two retained literature records, providing a particularly limited basis for answering the full panel.

The highest same-protein semantic agreement occurred for INSR (0.863 across 30 comparable questions), EGFR (0.847 across 33), and SRC (0.824 across 29). INSR and EGFR had no coverage mismatches. The lowest agreement occurred for CSNK1A1 (0.465 across 20 comparable questions), NEK1 (0.538 across 21), and GUCY2F (0.568 across 19). These proteins had seven, seven, and six coverage mismatches, respectively. Thus, lower aggregate agreement often accompanied differences in which questions the two sources answered. It cannot be attributed entirely to conflicting biological statements.

Question level agreement showed a related distinction. Gene symbols matched fully for all 50 proteins, while active kinase domain status and inactivating variants had mean self scores of 0.939 and 0.902 among comparable cases. The mean self scores for localization and substrates were 0.823 and 0.766, with 44 partial matches for each question. Tissue expression and cancer types had lower mean self scores of 0.406 and 0.302. This pattern indicates that shared broad biological knowledge can coexist with differences in lists of tissues, substrates, or disease contexts. The complete protein rankings and question summaries are provided in Supplementary Tables S1 and S2.

### Semantic scores distinguished matching proteins from alternative literature sets

The mean score for a UniProt answer set compared with its own literature answer set was 0.692, with a median of 0.703 and an interquartile range of 0.646–0.742 (Figure 2A). The mean across all 2,450 comparisons between different proteins was 0.287. For each UniProt query, we identified the highest score among its 49 alternative literature sets. These best alternative scores averaged 0.471, giving a mean self advantage of 0.221 (Figures 2B and Figure 3).

**Figure 2.**
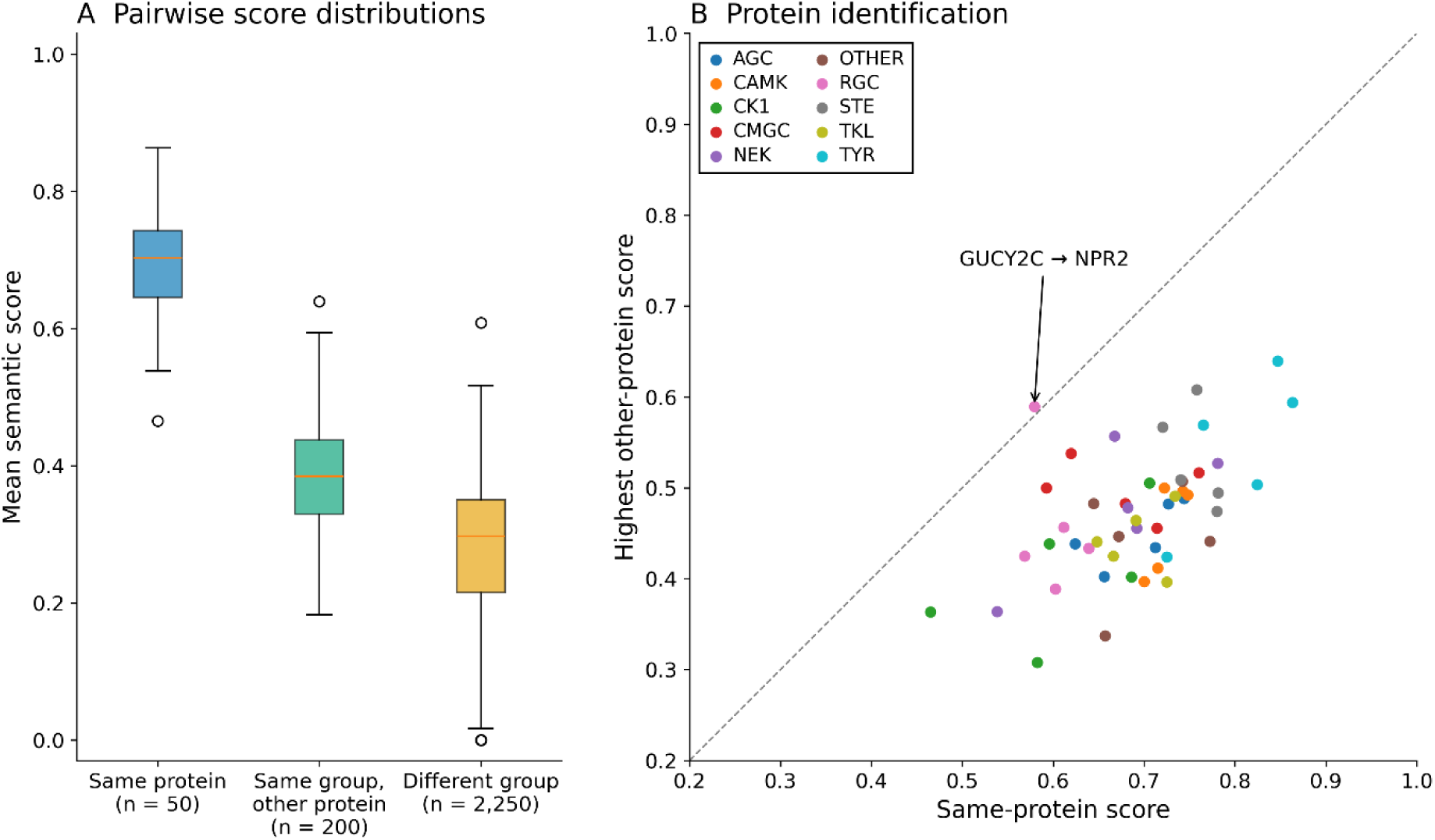
Score distributions and discrimination of matching proteins. **A)**. Boxplots showing scores for matching proteins, different proteins within the same group, and proteins from different groups. Boxes show the median and interquartile range; whiskers extend to observations within 1.5 interquartile ranges, with remaining observations shown individually. **B**) Comparison of each self score with its highest alternative literature score. Points below the identity line favor the matching protein. GUCY2C is the only point above the line. Comparisons share source answers, and the displayed observations are not independent replicates.

**Figure 3.**
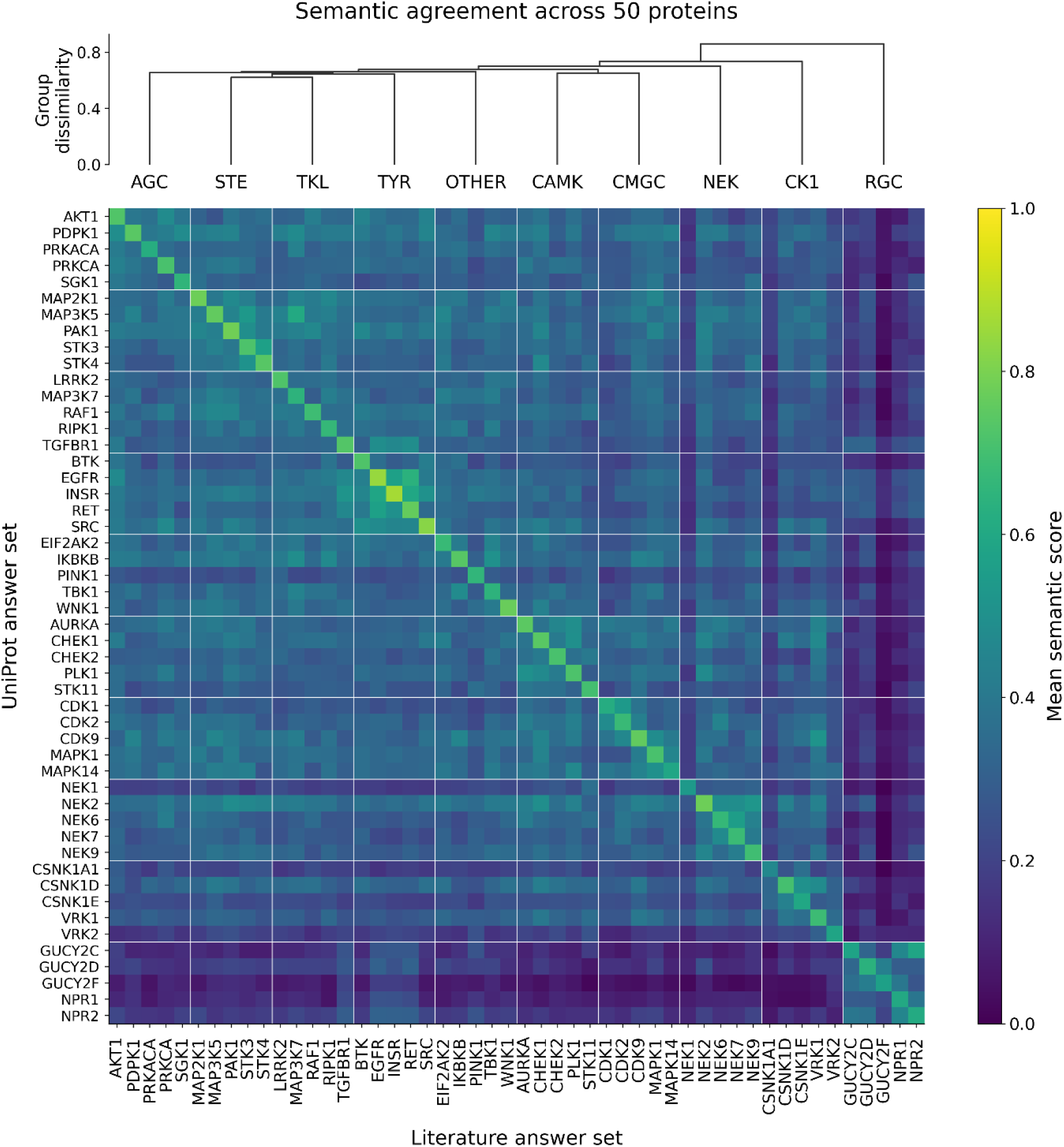
Semantic agreement between UniProt and literature answer sets. The matrix contains all 2,500 comparisons of 50 UniProt answer sets against 50 literature answer sets. The ten supplied kinase groups are ordered by average-linkage hierarchical clustering using one minus their mean between-group semantic score pooled across both source directions. The dendrogram shows the resulting group relationships; its heights indicate answer dissimilarity, not evolutionary distance or statistical significance. The same group order is used on both axes, with the five proteins in each group kept together and ordered alphabetically by gene symbol. White boundaries separate groups. Each cell is the mean of numeric scores across the 35 biological questions, with one-sided coverage gaps scored zero and jointly unreported questions excluded. Matching accessions form the diagonal. Symmetrization is used only for group clustering; the displayed matrix retains its original directed scores because each axis represents a different source.

The matching literature set ranked first for 49 of the 50 UniProt queries. This result was not solely attributable to the explicit identity question. Excluding gene symbol retained first place for the same 49 queries, and excluding both gene symbol and kinase family did so as well. In the latter sensitivity analysis, mean self agreement was 0.673 and mean agreement between different proteins was 0.306 (Supplementary Table S5). These analyses removed questions from the aggregation of existing judgments; they did not rerun the model with identifiers hidden.

Several of the closest alternative matches involved related proteins. CDK1 had a self score of 0.620 and a best alternative score of 0.538 against CDK2. MAPK14 scored 0.593 against itself and 0.500 against MAPK1, while NEK6 scored 0.667 against itself and 0.557 against NEK7. Such comparisons show that shared characteristics can produce appreciable similarity without eliminating the self advantage.

GUCY2C was the sole exception. Its UniProt answers scored 0.579 against GUCY2C literature and 0.589 against NPR2 literature, a difference of 0.010. The self comparison included 29 numeric questions and ten coverage mismatches, whereas the NPR2 comparison included 28 numeric questions and four coverage mismatches. Both GUCY2C and NPR2 belong to the RGC (receptor guanylyl cyclase) kinase group with inactive kinase domains. Shared receptor and guanylyl cyclase descriptions contributed to the alternative match. NPR2 literature also received full matches for signal peptide status, atypical kinase status, and activating variants, where the self comparison received coverage mismatches.

Inspection revealed that question availability and interpretation also affected the ranking. The GUCY2C UniProt answer to the question of proteolytic processing mentioned a precursor and chain boundaries while stating that processing was not reported. Against the GUCY2C literature answer, this question was not comparable; against the NPR2 answer, which mentioned a signal peptide cleavage site, it received a full match. Other questions, including inhibitors and autoinhibition, contributed zeros to the self comparison but were excluded from the NPR2 comparison when both answers were judged unreported. When the two aggregate scores were recomputed over the same 26 questions with numeric judgments in both comparisons, GUCY2C scored 0.646 against itself and 0.596 against NPR2. This case shows that the exception reflected shared biology together with reporting decisions and unequal denominators, rather than a consistent preference for the wrong protein across the same questions.

### Group assignments were reflected in answer similarity

Different proteins within the same kinase group had a mean score of 0.384 across 200 directed comparisons, compared with 0.279 across 2,250 comparisons between groups. Their median scores were 0.384 and 0.297, respectively, and their interquartile ranges were 0.329– 0.438 and 0.215–0.350, respectively. Within-group scores therefore tended to be higher, although the distributions overlapped. A group label permutation analysis preserved the paired protein identities and five proteins per group. None of 20,000 shuffled assignments produced a within-minus-between difference as large as the observed 0.106, giving a corrected Monte Carlo P value of 0.000050.

All ten groups had a higher within-group mean than their outgoing between-group mean. TYR and STE had the highest within-group means, 0.465 and 0.436, respectively. AGC had a mean of 0.388 within the group and 0.303 against other groups. OTHER and TKL had smaller differences of approximately 0.058. These group summaries were descriptive; the analysis did not establish pairwise significance for differences between individual group means.

The TYR kinase group also had the highest answer coverage, with 146 of 175 possible UniProt answers and 148 of 175 literature answers reported. CK1 had the lowest UniProt coverage, 94 of 175, whereas RGC had the lowest literature coverage, 99 of 175. Mean self agreement was highest for TYR (0.805) and STE (0.756) and lowest for RGC (0.600) and CK1 (0.607). These differences show that coverage and consistency can vary together, but they do not identify either one as the cause of the other. Group membership, selected proteins, publication availability, and the suitability of the questions are all potential contributors.

### Receptor guanylyl cyclases formed a distinct functional profile

RGC emerged as the most distinct of the ten groups in both the summary statistics and hierarchical clustering. Its within-group mean semantic agreement score was 0.350, compared with an outgoing mean of 0.136 against the other groups—the largest within-versus-between difference observed. When both directions of the RGC–non-RGC comparisons were pooled, the mean score was 0.139 across 450 comparisons. Consistent with these low cross-group scores, hierarchical clustering placed RGC on the outermost branch of the dendrogram, joining the other nine groups only at the highest level of group dissimilarity (Figure 2). The heatmap likewise showed a relatively dark RGC row and column outside the RGC block, indicating low semantic agreement between RGC proteins and members of the conventional kinase groups.

Question-level analysis identified especially strong RGC separation for substrates, transmembrane status, additional domains, and downstream pathways. Within RGC, the corresponding mean scores were 0.973, 1.000, 0.787, and 0.390, respectively, compared with 0.000, 0.127, 0.082, and 0.006 in comparable cross-group cases. Because each question-level mean was calculated from its own set of numeric judgments, these values are descriptive comparisons rather than a formal decomposition of the overall RGC group effect.

This pattern is consistent with the distinctive enzymology represented in the source answers. Receptor guanylyl cyclases contain membrane and cyclase components and produce cyclic GMP, while conventional kinase questions frequently concern protein phosphorylation.

Experimental studies established membrane guanylyl cyclases as natriuretic peptide receptors and identified GC-C as a receptor for heat-stable enterotoxin [21]. GUCY2F, the retinal cyclase RetGC-2, was characterized in photoreceptors, illustrating further specialization within this group [22]. Shared membrane topology, domain architecture, and cyclase activity therefore help explain the internal similarity of RGC proteins and their separation from the other groups, while distinct tissue and regulatory contexts limit agreement among RGC proteins themselves. The position of RGC as the most divergent group also highlights an ambiguity in applying the same question about substrates to a cyclase and a protein kinase.

## Discussion

This study found substantial complementarity between answers derived from UniProt records and answers derived from their cited literature. Most reported answers were shared at the level of coverage, but each source also supported questions unanswered by the other. Literature had a modest overall coverage advantage, concentrated most clearly in autoinhibition and inhibitor types after correction for multiple comparisons. UniProt showed descriptive advantages for tissue expression, cancer types, and processing. These results support examining source strengths at the level of individual questions rather than assigning a single measure of superiority to either source.

The restriction to UniProt cited publications is central to that interpretation. The literature arm was not an unrestricted search for all available knowledge about each protein. It reexamined publications already connected to the corresponding UniProt record. Additional reported answers may therefore reflect differences in what the two representations preserve, the scope of individual annotations, or the extraction procedure. They cannot be interpreted automatically as newly discovered knowledge or as omissions by database curators. Furthermore, both answer sets were generated by the same model family, and the literature evidence included target metadata derived from UniProt. The two processing streams were separate, but their information sources were not fully independent.

The high rate of correct self ranking provides evidence that the answer profiles retained protein specificity. A generic description shared across many signaling proteins would not be expected to distinguish nearly every matching literature set from 49 alternatives. The persistence of 49 correct first rankings after removing gene symbol and family questions strengthens this observation. However, this remains a test of discrimination within a selected panel. Related targets outside the panel might be harder alternatives, and the model saw accession labels and protein names within answer texts. The result therefore does not constitute a blinded protein identification benchmark or an estimate of biological answer accuracy.

The GUCY2C exception provides a useful limit on what a single aggregate score can represent. The mean comparison score jointly reflects semantic overlap and reporting completeness, because a coverage mismatch contributes zero to the score. This choice prevents one-sided missing information from being treated as agreement. It also means that a literature set containing additional valid findings can receive a lower score when its UniProt counterpart is silent. Excluding questions that are unreported in both sources introduces a second effect: different candidate pairs can be averaged over different questions. In the GUCY2C comparison, restricting the analysis to common numeric questions restored the self advantage. This does not justify changing the primary metric to force the expected ranking; it identifies the denominator as a source of sensitivity that should accompany interpretation of the score.

Several other features of the question panel affect interpretation. Some questions ask for specific entities or mechanisms, whereas others ask only whether a property exists. Two unrelated proteins can both have inactivating variants and correctly receive matching yes answers despite having different variants. Conversely, two accurate substrate lists may overlap only partly because the underlying experiments examined different targets. Equal weighting treats these different forms of information as interchangeable contributions to one score. Closely connected questions about regulators, activation, modifications, pathways, and cellular consequences can also count related evidence more than once. The observed separation between and within groups is consequently a property of this question panel and scoring scheme, not a direct estimate of sequence similarity or evolutionary distance.

Limited coverage also has more than one explanation. Secretion, receptor status, and signal peptides are not routinely discussed for every intracellular signaling protein. Drug resistance and cancer causality depend on particular experimental or clinical contexts. Terms such as inactive conformation, catalytic inactivity, pseudokinase, and autoinhibition refer to different concepts, although they may appear in similar sentences. The RGC comparisons make this problem especially visible because a catalytically active cyclase can contain a kinase homology domain without established protein kinase activity. An answer can be informative about one of these properties while remaining insufficient for another.

Answer coverage indicates how much information was recovered but does not fully describe what is available in the sources. A reported answer may be a positive claim, a negative result, or a qualified observation. An answer marked as not reported may mean that the evidence is absent, was missed during extraction, or was excluded by a strict interpretation of the question. The question may also be unsuitable for the protein. Classifying answers only as reported or not reported does not distinguish these cases. Additional categories could separate explicit negative evidence, uncertainty, and questions that do not apply, provided each category is clearly defined and supported by the source.

The extraction and evaluation steps also have limitations. Filtering literature records for the target protein early in the analysis helps prevent findings about an interaction partner or paralog from being assigned to the target. However, this step may exclude relevant records if protein names or other identifying information were incompletely extracted. Extracting text from full articles, processing it in chunks, and combining the results can also lose details or caveats. Retained records may include passages describing the broader experimental context, so filtering does not ensure that every statement used to generate an answer refers specifically to the target protein.

Statistical uncertainty must also be interpreted at the appropriate level. The 87,500 question comparisons are not independent biological observations. They repeatedly use the same proteins, source answers, and overlapping publications. Our resampling analyses therefore retained protein or group structure rather than treating each matrix cell as an independent replicate. Nevertheless, only five proteins represented each supplied group, and selection was not established as random. The question tests were exploratory, protein relationships may introduce residual dependence, and the bootstrap intervals do not include uncertainty from rerunning the language models. The statistical findings are conditional on this collection and its saved outputs.

A subsequent validation study should compare extracted claims against expert assessments linked to specific passages and experimental contexts. Such an evaluation could measure unsupported claims and missed evidence separately and assess whether results persist across models and repeated runs. Testing the scoring rubric with controlled differences in specificity, negation, and missing information would help distinguish semantic judgment errors from genuine source differences. Within those limits, the present work provides a reproducible way to locate complementary evidence, characterize coverage gaps, and identify comparisons that warrant closer biological review.

## Materials and Methods

### Protein panel and group assignments

The analysis comprised 50 proteins selected from the sequence collection of the 484 human protein kinases associated with Modi and Dunbrack (2019). Five proteins with the highest citation counts in UniProt were selected from each of the ten kinase groups: AGC, CAMK, CK1, CMGC, NEK, OTHER, RGC, STE, TKL, and TYR. The panel included conventional protein kinases as well as five receptor guanylate cyclases in the RGC group (GUCY2C, GUCY2D, GUCY2F, NPR1, and NPR2), which were analyzed as related signaling proteins containing catalytically inactive kinase-homology domains. The selected accessions and their group assignments are provided in Supplementary Tables S2 and S3.

### Literature scope and article extraction

For selected protein kinases, we retrieved publications cited in UniProt and downloaded the corresponding articles from PubMed Central (PMC) [23], Europe PMC [24], and journal websites. Retracted publications were identified based on PubMed metadata and excluded from further analysis. The retrieved articles were available in multiple formats, including PDF, JATS/NLM XML, publisher-specific XML, and HTML. PDF files were converted to TEI XML using GROBID. Articles in all formats were subsequently converted to plain text, with the bibliography sections removed, and the resulting text was concatenated for downstream analysis. Information extraction was performed using GPT-5.6 Luna with high reasoning effort. The complete prompt used for information extraction is provided in the Supplementary Materials.

### Target identity filtering of literature

An AI-assisted canonical target map was constructed from the accession, gene name, synonyms, and full and short protein names in the local UniProt records. Using ChatGPT (AI), literature entities were matched against accessions, normalized gene symbols and protein names, aliases, and curated historical names. AI-assisted normalization accounted for capitalization, punctuation, Greek-letter variants, and selected naming conventions. Generic kinase labels and unresolved family mentions were considered insufficient by the AI filtering procedure for target assignment. The AI procedure combined general matching rules with a small, curated alias dictionary and used language-model-assisted adjudication of target identity.

Records were retained through AI-assisted filtering when a matched entity represented a primary, co-primary, or contextual target. Claims linked to the AI-matched target were retained together with entities required to represent those claims, while associated features, variants, and modulators were restricted using their entity relationships. Records that remained unresolved after AI assessment were quarantined. Primary and contextual evidence identified through AI filtering was combined for answer generation, with unresolved categories excluded. The original records were preserved, and AI-assisted filtering decisions were recorded in an audit table. The manual-review category referred to identity resolution and did not designate review articles as a publication type.

### Question answering

Both sources were evaluated using the same AI-assisted workflow and the same predefined 35 questions. The complete wording is provided in Supplementary Table S1. The questions covered identity and classification, catalytic and regulatory features, topology and localization, substrates and signaling, modifications, variants, inhibitors, and disease associations. Each AI-generated answer included a text response, a reporting state, and evidence excerpts. “Reported” indicated that the AI extraction supplied a claim relevant to the question; “not reported” indicated that supporting information was not identified by the AI workflow. Absence of evidence in the AI-generated output was not intended to imply that a biological feature was absent.

Answers were generated by an AI model, GPT-5.2, through the OpenAI Responses endpoint and Batch workflow. The UniProt input provided to the AI model was the accession-specific flat file, including its annotation and feature text. The literature input provided to the AI model consisted of the retained full-article extractions for the same target. The AI model was instructed to use the supplied evidence, preserve uncertainty and experimental qualifications, and distinguish the target from constructs, interacting proteins, and downstream readouts.

Literature extractions were divided at complete record boundaries into chunks targeted to a maximum of 450,000 characters for AI processing. The final retained collection produced 192 AI extraction requests. Candidate AI-generated answers from these chunks were synthesized by the AI model in 50 further requests, yielding one answer set per protein. The standard AI answer-generation request allowed 10,000 output tokens; two incomplete AI extraction requests were retried with 20,000. No explicit reasoning-effort parameter was supplied for AI answer generation. AI outputs were returned in a constrained structured format and normalized to the common question list. Final coverage analyses used the saved AI answer-generation states, separately from states subsequently assigned by the AI-based semantic adjudicator.

### Semantic evaluation and adjudication

All UniProt answer sets were compared against all literature answer sets using an AI-based semantic evaluation. The primary AI evaluation used GPT-5.6 Sol with reasoning effort set to high and a maximum of 20,000 output tokens. Each AI evaluation request contained all 35 questions for one protein pair and the two AI-generated answer texts with their reporting states. The AI evaluator was instructed to compare meaning using the supplied answers and not to infer biology from accessions or external knowledge. Source evidence excerpts and original articles were not supplied to the AI judge. Accession labels were present, so the AI evaluation was not formally blinded.

The AI evaluation rubric classified each comparison into one of five categories: (1) **semantic match**, in which the two reported answers conveyed equivalent meaning and received a score of 1.0; (2) **partial match**, in which the answers showed meaningful semantic overlap but differed materially in specificity, scope, qualification, or other content and received a score of 0.5–0.9; (3) **mismatch**, in which both answers were reported but their meanings were materially different or contradictory and received a score of 0.0–0.4; (4) **coverage mismatch**, in which exactly one answer was reported and the other was unreported, resulting in a score of zero; and (5) **not comparable**, in which both answers were unreported and the comparison received a score of zero but was distinguished from a mismatch because neither source provided information for semantic comparison. Explicit negative statements and qualified claims could be classified by the AI evaluator as reported information and were compared using the same semantic criteria.

### Score summaries and sensitivity analyses

For each UniProt–literature pair, the aggregate score was the arithmetic mean of all numeric question scores. Every comparable question received equal weight. Coverage mismatches therefore remained in the denominator with score zero, whereas questions classified as not comparable were omitted. The number of comparable questions and coverage mismatches was retained for every pair. Because the two axes represented different source types, the matrix was directed and was not assumed to be symmetric.

For each UniProt query, its self score was compared with the highest score among the 49 other literature sets. Within-group comparisons excluded the same accession and comprised 200 directed pairs, with 20 per group. Between-group comparisons comprised 2,250 directed pairs. Group specific between-group means treated the group of the UniProt query as the reference. The RGC question analysis additionally pooled both directions of comparisons involving one RGC and one non-RGC protein. Sensitivity analyses excluded the gene symbol question, excluded both gene symbol and kinase family, and restricted the GUCY2C case study to questions with numeric scores in both competing comparisons.

### Statistical analysis

Answer coverage was the proportion of question entries labeled reported in each source. For individual questions, the paired comparison counted proteins with a UniProt-only answer and proteins with a literature-only answer. Two-sided exact McNemar tests were calculated as binomial tail probabilities conditional on the number of discordant pairs [25]. Holm adjustment controlled multiplicity across all 35 question tests, with adjusted P values below 0.05 treated as exploratory evidence of a coverage difference [26].

For the overall coverage difference, 20,000 bootstrap samples resampled five proteins with replacement within each of the ten groups. Both sources and all questions for each sampled protein were kept together. The 2.5th and 97.5th percentiles defined the reported 95% confidence interval. A separate exact group-level permutation enumerated all 1,024 assignments obtained by either retaining or swapping the source labels jointly for all proteins and questions in each group. Its two-sided P value was the proportion of assignments with an absolute coverage difference at least as large as observed.

Group structure in the score matrix was assessed by 20,000 random permutations of group labels across 50 proteins, maintaining five labels per group. The same permuted label followed a protein on both axes. The statistic was the within-group mean minus the between-group mean, excluding self comparisons. The one-sided Monte Carlo P value was calculated as the number of permuted statistics at least as large as observed, plus one, divided by 20,001. These analyses assumed exchangeability under their respective null hypotheses and were interpreted conditionally on the selected panel. The random seed was 20260910. Statistical calculations and figures used Python and NumPy, with Matplotlib for visualization.

## Supplementary materials

Supplementary Table S1. contains the complete question panel, coverage counts, exact paired tests, Holm adjusted P values, and same-protein semantic summaries. Supplementary Table S2 contains all 50 proteins, group assignments, retained literature counts, source coverage, self scores, and best alternative matches. Supplementary Table S3 summarizes coverage and scores by group. Supplementary Table S4 contains directed group-pair score distributions with self comparisons excluded. Supplementary Table S5 reports the question exclusion sensitivity analyses. Supplementary Table S6 contains all 35 questions, answer texts, and competing judgments for GUCY2C and NPR2. Supplementary Table S7 reports question level RGC comparisons.

## Data availability

The data generated in this study are available in supplementary materials.

## Acknowledgements

We would like to thank Drs. Lisa Kinch and R. Dustin Schaeffer for helpful discussion and suggestions. Qian Cong is a Southwestern Medical Foundation scholar. We thank TACC for providing computing resources. This research is supported by National Institute of General Medical Sciences (1R35GM160468-01) to Q.C., the Welch Foundation grant V-I-0004-20230731 to Q.C., National Institute of Allergy and Infectious Diseases (1K99AI180984-01) to J.Z., the CPRIT grant RR260055 to J.Z., National Institutes of Health (GM127390) to N.V.G., and the Welch Foundation grant I-1505 to N.V.G.

## Conflict Of Interest Statement

The authors declare no competing interests regarding the content of this study.

## Notes

### Competing Interest Statement

The authors have declared no competing interest.

